# Presence Hallucination Induction through Robotically Mediated Somatomotor Conflicts – a pooled analysis of 26 experiments

**DOI:** 10.1101/2025.09.26.678742

**Authors:** Herberto Dhanis, Jevita Potheegadoo, Nathan Faivre, Masayuki Hara, Fosco Bernasconi, Olaf Blanke

## Abstract

Hallucinations are significant symptoms in psychiatric and neurodegenerative diseases, that may indicate advanced disease progression or worse disease forms. They are also frequent in healthy individuals, especially elderly or bereaved. Despite their relevance, studying hallucinations in controlled laboratory conditions remains challenging given the limited procedures inducing clinically relevant hallucinations. We have previously developed a robotics-based protocol capable of inducing a specific clinically relevant hallucination, presence hallucination (PH), in both healthy individuals and patients. Using this approach, we have systematically investigated the sensitivity and intensity of PH-induction, as well as its effects on various sensory, behavioral and cognitive aspects. In the present study, we pooled individual-participant data from 26 in-house experiments (totaling 580 individuals) and conducted a Bayesian analysis to estimate effects and moderators of PH-induction. PH-induction was reliably induced with a medium effect-size, and individuals with schizotypal traits were more sensitive. Furthermore, we identified that PH-induction may not strongly depend on altered agency, but found a synergistic relationship between passivity and induced PH. Collectively these results elucidate the role of somatomotor processes in aberrant own body perceptions, advance understanding of psychosis, and provide powerful statistical priors for future studies.

## 1 Introduction

Hallucinations are multifaceted and heterogeneous phenomena in which an individual experiences abnormal perceptual sensations in the absence of corresponding external stimuli (American Psychiatric Association, 2013). The clinical relevance of hallucinations has been demonstrated trans-diagnostically, in psychiatric conditions such as schizophrenia (Whitfield-Gabrieli and Ford, 2012), in neurological disorders such as stroke (De Haan et al., 2007; Flint et al., 2005), and neurodegenerative diseases like dementia with Lewy bodies (Nagahama et al., 2007) and Parkinson’s disease (Ffytche et al., 2017), but also in healthy individuals (e.g., Badcock et al., 2020).

Although most studies investigate visual and auditory hallucinations (e.g., Corlett et al., 2019; Honcamp et al., 2022; Zarkali et al., 2022), other sensory systems may also be affected and give rise to non-visual and non-auditory hallucinations, for example: vestibular or somatosensory hallucinations (Blom and Sommer, 2012; Cheyne and Girard, 2009). In cases such as autoscopic phenomena, broad systems involved in multisensory perceptual body representations may be implicated and individuals experience hallucinations of their own body (e.g., autoscopic hallucinations, heautoscopy, and out-of-body experiences; (Blanke et al., 2004; Blanke and Mohr, 2005; Blondiaux et al., 2021; Brugger et al., 2006). One of such hallucinations is the Presence Hallucination (PH): the convincing and realistic sensation of feeling someone close by when no one is actually there (Ajuriaguerra and Hecaen, 1952; Brugger et al., 1997, 1996; Critchley, 1955). PH is most often experienced behind or to the side of the afflicted individual, sometimes sharing the same body traits, and mimicking movements, postures and actions (Blanke et al., 2008, 2003; Brugger et al., 1997, 1996; Potheegadoo et al., 2022). Despite its high degree of perceived realism, PH has no clear sensory components, as opposed to formed visual or auditory-verbal hallucinations (Brugger et al., 1997; Picard, 2010). PH is frequent in schizophrenia (46%, Llorca et al., 2016), dementia with Lewy bodies (20%, Nagahama et al., 2010), and to a lesser degree has also been described in epilepsy (Blanke et al., 2014; Picard, 2010) and in healthy individuals undergoing extreme circumstances such as high-altitude mountaineering (Messner, 2016; Brugger et al., 1999), shipwrecks (Critchley, 1943; Geiger, 2009) or bereavement (Kamp et al., 2022; Ratcliffe, 2021). In Parkinson’s disease, PH is particularly relevant as it is one of the most common and early hallucinations experienced (Fénelon et al., 2011a), with around 42% of patients experiencing PH before diagnosis and approximately 15% before the onset of motor symptoms (Pagonabarraga et al., 2016). Moreover, the early occurrence of PH in these patients has been associated with other psychiatric symptoms (Ffytche et al., 2017; Lenka et al., 2019) as well as faster cognitive decline (Bejr-kasem et al., 2021; Bernasconi et al., 2023).

Several models have been put forward to account for the occurrence of PH, offering different explanations for this phenomenon (Barnby et al., 2023a; Blanke et al., 2014; Fénelon et al., 2011b). Neurological evidence suggests that disrupting the integration of proprioceptive, tactile, and motor signals in multisensory and somatomotor integration areas such as the temporoparietal junction may lead to PH (in stroke: Blanke et al., 2003; in invasive electrical stimulation of an epileptic patient undergoing pre-surgical evaluations: Arzy et al., 2006; in neurological patients more generally: Blanke et al., 2014). This somatomotor mismatch hypothesis suggests that the discrepancy between actual sensory signals and expected sensory feedback may trigger the experience of PH, due to altered proprioception or motor control. Moreover, PH symptomatology, neurological evidence, and some patients’ experiences of affinity and self-identification with the felt presence, has led to grouping PH with alterations of the body schema and altered somatomotor processing, and ultimately to consider PH as misperceived double of the self-arising due to impaired somatomotor processing (Barnby et al., 2023a; Case et al., 2020). PH has also been shown to be triggered by neuro-chemical or pharmacological alterations (Dupuis, 2022; Somer et al., 2023; Wolff et al., 2019) though the exact mechanisms remain unclear. For instance, medications used for treating motor symptoms in Parkinson’s disease, such as dopaminergic agonists and levodopa that affect dopamine levels can enhance hallucinations. PH might be a mild form of such hallucinations induced or exacerbated by dopaminergic treatments. Besides dopamine, serotonin-glutamate interactions and acetylcholine deficits, along with brain region dysfunction (e.g., in the temporoparietal junction), could be central contributors (Ballard et al., 2013; Jalal, 2018). Other authors have proposed PH as a form of "social" hallucination due to their strong connection to autobiographical and affective memory networks (Barnby et al., 2023b; Fénelon et al., 2011b; Kamp et al., 2019). PH, as experienced in Parkinson’s disease, in bereavement, or during sensory deprivation and stressful situations, often involve the sensation of a familiar person, and the feeling of presence is frequently associated with emotionally significant individuals, such as a deceased close other (e.g., spouse, sibling; Ratcliffe, 2021; Steffen and Coyle, 2011). This suggests that PH may arise from the activation of autobiographical memories, particularly when the brain reconstructs the presence of a person based on past experiences, often in the absence of perceptual clues. Misattribution of these internal reconstructions to an external source can create the subjective feeling of an actual presence. Additionally, PH can be influenced by prior knowledge and perceptual expectations, further linking it to social cognition (Barnby et al., 2023b). This involvement of both autobiographical memory and social networks supports the idea that PH is shaped by emotional and relational contexts, reinforcing its classification as a "social" hallucination. Despite these different hypotheses and despite the clinical relevance of PH and available data, precise descriptions of its mechanisms remain elusive due to a variety of factors. These models might not be mutually exclusive, and the experience of PH is likely multifactorial, involving a combination of neurological and psychosocial factors (Alderson-Day et al., 2023; Corlett et al., 2019).

The occurrence of hallucinations such as PH is generally unpredictable, happening during daily life activities and most often far from clinical settings. Prejudices and stigma associated with hallucinatory experiences also refrain many patients (and healthy individuals) from reporting them (Pagonabarraga et al., 2014; Wood et al., 2015), further hindering their investigation. In Parkinson’s disease, this has been further exacerbated by the absence of a systematic screening by clinicians (Chan and Rossor, 2002). Even when hallucinations are reported, their recollection and interpretation is likely confounded by several patient and clinician biases (Berrios and Marková, 2002; Nisbett and Wilson, 1977; Ravina et al., 2007). Moreover, the absence of clear sensory components in PH (as present in visual, auditory, or tactile hallucinations) has made their classification and scientific testing difficult, likely giving rise to the large range of proposed mechanisms. In that sense, experimental procedures that could induce PH, without any disease, without any alteration of consciousness, without pharmacological interventions (Baggott et al., 2010; Timmermann et al., 2018), and in real-time controlled laboratorial conditions, would be highly beneficial to better understand the phenomenology of these hallucinations.

Recently, we have developed a new procedure to overcome those limitations. Integrating clinical observations of impaired somatomotor processing in PH (Arzy et al., 2006; Blanke et al., 2014, 2003) with advances in cognitive neuroscience and robotics, we designed a protocol capable of inducing PH in fully controlled experimental conditions, thus allowing a real-time assessment of the hallucinations (Bernasconi et al., 2022). Through this protocol, individuals are exposed to specific spatio-temporal conflicts between motor, tactile and proprioceptive cues from a forward extended arm and from a tactile cue applied on the back, triggering a robot-induced PH (riPH). Importantly, other controlled somatomotor stimulations using the same robotic device do not lead to riPH. This induction of PH is accompanied by modulations of other bodily experiences, notably passivity experiences (PE; robot-induced PE: riPE), the sensation that someone else is applying the touch on the participant’s back (Mlakar et al., 1994) as well as decreases in sense of agency, the sensation of being in control of one’s actions (Haggard and Chambon, 2012). To date this riPH protocol has been replicated in multiple experiments and has also been used to study the impact of riPH in different perceptual and cognitive processes, such as auditory-verbal processing, auditory-verbal hallucinations (Salomon et al., 2020), auditory misperceptions (Orepic et al., 2021), fluency and memory tasks (Serino et al., 2021), metacognition (Faivre et al., 2020), and a numerosity estimation task in VR (Albert et al., 2024). The robotic procedure has also been extended to MRI (Bernasconi et al., 2021), allowing the investigation of the neural mechanisms of riPH (Bernasconi et al., 2021; Dhanis et al., 2024) and other associated experiences such as riPE (Dhanis et al., 2022).

Given the large number of different studies, experimental settings and research questions we have applied this protocol to, including ongoing translation to clinical populations (Bernasconi et al., 2021; Salomon et al., 2020), we aimed to quantify the effect size of riPH, identify population-traits that make individuals more sensitive to the procedure, and determine the optimal setup by investigating various within- and between-experiment parameters. To this aim, we carried out a Bayesian estimation analysis on pooled individual-participant data from 26 experiments performed in our laboratory, both published and pilot data to avoid publication-bias effects, which induced riPH in a total of 580 healthy individuals. In doing so, we estimated that riPH can be reliably induced in the general population with a medium effect size and that individuals with delusional ideations are more prone to experiencing riPH. Additionally, we propose that sensations which typically accompany riPH, such as riPE and changes in the sense of agency (Bernasconi et al., 2021; Blanke et al., 2014), but that have distinct neural mechanisms (Dhanis et al., 2022), distinctly affect riPH.

## 2 Methods

### 2.1 Studies and Data

The studies included here were conducted in the Laboratory of Cognitive Neuroscience and are either published or constitute pilot data which we are releasing in an effort to counter the publication bias that notoriously affects large analyses that pool data from multiple studies (Dickersin, 2005; Sharpe, 1997). Supplementary Table S1 shows the included studies and pilots.

All included studies were approved by the Swiss ethics committee and written informed consents were obtained from all participants prior to their inclusion to each study.

### 2.2 Participant’s demographics and trait characteristics

We investigated whether participants’ trait characteristics influenced the induction of riPH and other subjective experiences. This included their age, handedness score as assessed by the Edinburgh Handedness Inventory (EHI) (Oldfield, 1971), and delusional ideation score as assessed by the Peter’s Delusions Inventory (PDI) (Peters et al., 2004). Sex at birth was included as a covariate of no-interest.

### 2.3 General Experimental Setup

All the experiments included in this analysis used the same base protocol (Bernasconi et al., 2022). Participants were blindfolded with an eye-mask and white noise was presented through headphones to isolate them from surrounding noise. While isolated, participants had a handle attached to their right index finger and moved the front part of a robotic system (front-robot; Figures 1A-B) by performing forward and backward poking movements. At the same time, these movements were transmitted and reproduced by a back-robot (Figures 1A-B), onto the participant’s backs (Figures 1C-D). When operating this system with the two robots in synchrony participants do not experience riPH, but rather report control sensations, such as self-touch. It is only when a temporal conflict is introduced between the participants’ movements and the tactile feedback on the back (asynchronous condition), thus increasing the perceived somatomotor conflicts, that participants report riPH. The change between synchronous and asynchronous conditions is noted in *Equation 1* as Asynchrony. To assess the subjective experience induced by the somatomotor robotic stimulation, each participant was asked to respond to a questionnaire immediately after each session (i.e. a session comprises the duration of manipulation under the same condition). For each questionnaire item, participants had to rate on a 7-point Likert’s scale, the intensity of the subjective experience (from 0 being “not at all”, to 6, being “very strongly”). The following two items were included in every experiment: “I felt as if someone was standing close to me (next to me or behind me)”, to assess the illusory sensation of riPH, and “I felt as if someone else was touching my body”, to assess riPE. Questions assessing other subjective sensations (i.e., agency, the experience of self-touch, anxiety) and different control questions were also pooled from the different studies (Supplementary Table S2).

**Figure 1.** is cut due to copyright but can be seen at Bernasconi et al. 2022 Nature Protocols.

### 2.4 Experimental parameters

Since the original PH induction study (Blanke et al., 2014), the various follow-up studies we have done included some differences in experimental parameters, for example, to adapt the paradigm to MRI settings, explore different control conditions, or investigate how riPH could affect behavioral or cognitive functions. Here, we describe all parameters that varied within and between experiments and that we aim to study whether they change the perception of riPH.

#### Within experiment

Every experiment repeated the period of robot manipulation at least twice, having one experimental condition with the front- and back-robot working in synchrony (participants received feedback on their back in synchrony with their movements), and another period where the robots moved asynchronously (500 ms of delay between the movements in the front and the feedback on the back). The order of these two experimental conditions was randomized across participants and this was included in our analysis (order). Some experiments included one additional condition, where the tactile feedback from the back-robot was given to the back of the hand rather than the back (hand comparator: back as usual or using hand for comparison), to investigate whether trunk-centered or peripheral stimulation influenced riPH (e.g. Faivre et al., 2020). The hand was placed in front of the participants to receive tactile stimulation, and an additional question measured riPH in the front as well.

#### Between experiments

Due to differences in experimental setup specific to each study, several parameters varied across the experiments. These included the duration of the somatomotor robot manipulation period (duration), the body-position given that participants were either standing (classical robot; Blanke et al., 2014) or supine (MR-compatible version; Bernasconi et al., 2021), previous exposure to the robotic stimulation (i.e., extensive use of the robot before the assessment of riPH), and the presence of concomitant tasks during the induction of PH (cognitive load). Cognitive load is encoded as a binary value (Yes/No), however, note that this simplification implies that we considered any concomitant task as cognitive load but the actual cognitive load might have varied between experiments. Supplementary Table S3 shows how these parameters varied across experiments.

### 2.5 Statistical Analysis

We conducted a Bayesian Estimation analysis to assess the effect of asynchrony and different experimental parameters, on the absolute ratings of riPH and accompanying sensations, measured at the end of each manipulation session. We selected a Bayesian framework for its ability to provide full distributions on the posteriors and followed literature recommendations (Kruschke and Liddell, 2018a, 2018b; Liddell and Kruschke, 2018). All analyses were performed with the use of the brms package (Bürkner, 2017) available for R (version 4.0.5).

To estimate the effect of the experimental parameters and age on the induced illusions we first fitted a full ordinal model on the data (*experiment model*), with a cumulative probit link-function with a flexible threshold to account for the fact that ratings were collected on an ordinal scale. The model equation was as seen below:

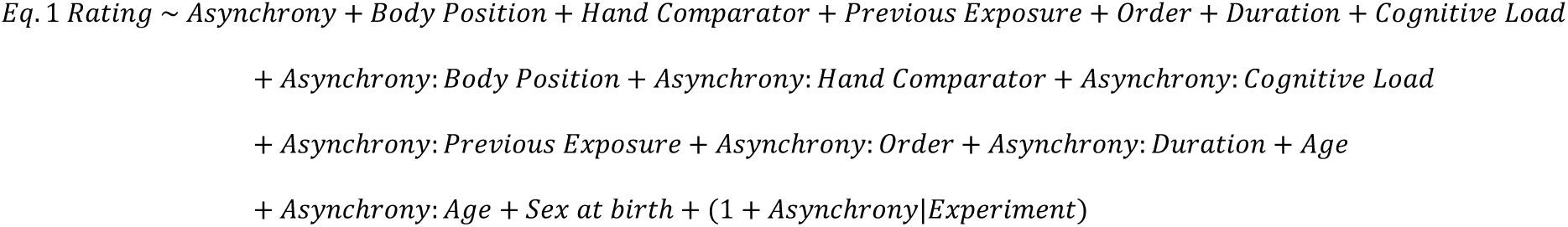

The inclusion of the interactions between asynchrony and body position, hand comparator, cognitive load, previous exposure, order and age respectively, question whether the latter parameters influence the main modulator of riPH (asynchrony). A random intercept term for experiment was also included, to capture potential variability not expressed in the model. Regarding model estimation, our model used a weakly informative Gaussian prior for the population-level predictors, 𝒩(0,5) and a weakly informative t-student prior for the group-level random effects, *t*_3_(0,2), as to regularize estimates without imposing strong assumptions.

Two more models were fitted separately in smaller populations to account for the effects of handedness and PDI scores, respectively. The models were the same as the experiment model but included added terms for handedness and its interaction with asynchrony (*handedness model*), or the same but applied to PDI (*schizotypy model*). These had to be assessed separately given that EHI and PDI were not available for the same population subsets, but to guarantee that these models still accurately preserved the variance of the experimental parameters, we used the estimates obtained from the *experiment model* as priors here for the experimental parameters. This allowed us to accurately maintain the modelling of variance brought up by the experimental parameters (as it had been assessed in larger pools of participants) and improve the estimates of EHI and PDI respectively. Note that we did not re-evaluate any experimental parameters here, as the goal was only to preserve their variance.

The beta estimates obtained from our models refer directly to effect size, or in other words are expressed in standard deviation (SD). A beta estimate of 0.5 for a parameter would mean that changing that parameter produces a change of half a SD in the ratings, being that this SD is the SD of the whole acquired data including all levels of all factors, for the question being analyzed. Importantly, this is with respect to the baseline of all other experimental parameters since our parameters often code absence vs use (for the asynchrony parameter, synchrony is baseline; age, EHI and PDI are centered). For each of the beta estimates we computed the 89% high-density intervals (HDI), as these are more stable to changes in effective sample size than their 95% counterparts especially when not in the upper thousands of samples (Kruschke, 2014). Once these were estimated, we used the Region of Practical Equivalence (ROPE) procedure to establish which experimental parameters modulated the ratings of each illusion.

In sum, this statistical procedure establishes a region around the null value, in this case 0, which is considered to be practically equivalent to the null value (Kruschke and Liddell, 2018a). We considered this ROPE to extend ± 0.1 standard deviations from the null value, which is equivalent to considering very small effect sizes as no practical effect. If the HDI of an estimate fell completely within this ROPE, then we considered this parameter to be practically equivalent to the null value, and hence not modulating the ratings. Otherwise, if the HDI fell completely outside the ROPE, we considered that the parameter had an effect different than the null value and hence modulating the experience of the induced sensation. If the estimate fell outside the ROPE, but its HDI overlapped with the ROPE, we could not explicitly conclude whether that parameter modulated the experience of the induced sensation differently than the null value. We also provide the percentage of how many posterior estimates fall inside the ROPE (this is the equivalent of comparing a 100% HDI to the ROPE). Finally, diagnostic plots can be found at the end of the supplementary information, in the section “Diagnostic Plots”.

### 2.6 Mediation Analysis for the different induced sensations

To assess potential relationships between riPH and accompanying sensations of riPE and agency, we modelled how sensitivity to changes in agency, and to riPH or riPE, affected the intensity of riPE or riPH, respectively. While sensitivity to induction refers to the induction of these illusions regardless of their intensity (i.e., any positive score difference between asynchronous and synchronous conditions), intensity of induction refers to the actual difference between the scores in the asynchronous versus synchronous conditions. We did so using an approach akin to mediation analysis, with special considerations. A typical mediation analysis would assess, for example, whether asynchrony modulates riPH, and whether further changes in agency further modulate riPH. We were additionally interested in investigating whether sensitivity to riPE would also modulate riPH intensity. Knowing that this variable is not independent from sensitivity to changes in agency added an additional problem (discussed here: VanderWeele and Vansteelandt, 2014). To circumvent this problem, we modelled a three-way interaction in stages between asynchrony, sensitivity to changes in agency, and sensitivity to riPH or riPE (depending on if the dependent variable was intensity of riPE or riPH. This approach ensures that we modulate any potential interaction between these variables, but included some interactions of no interest. The most strict way to model this question would be using Structural Equation modelling (as done here: Shelleby et al., 2014), however this is not possible with the type of data at hand.

## 3 Results

### 3.1 Included studies, characteristics and assessed parameters

Included in this pooled analysis are 580 healthy individuals (291 males) with a mean age of 24.1 years (SD: 4.9 years; range: 7 to 56 years), collected over 26 in-house experiments from 14 different studies. Of these experiments, 17 are published (Albert et al., 2024; Bernasconi et al., 2021; Blanke et al., 2014; Dhanis et al., 2024; Orepic et al., 2024, 2021; Salomon et al., 2020; Serino et al., 2021) and nine are pilot data that are being released now to provide accurate estimates beyond publication-bias (Supplementary Table S1). It should be noted that no participant was ever repeated in different experiments, as having participated in a previous experiment was an exclusion criterion of all experiments.

### 3.2 Effect of asynchrony of the somatomotor stimulation on robot induced PH

The effect of the robotically-mediated somatomotor stimulation on riPH (i.e. asynchrony) had positive but varied effect sizes across past experiments (Figure 2A). Hence it was one of the main goals of this pooled analysis to quantify its true effect. Our model provides evidence in favor of the asynchronous somatomotor stimulation leading to higher ratings of riPH than synchronous stimulation, with the standardized mode for its effect size set at 0.35 SD (HDI: [0.19, 0.48]; Figure 2B).

**Figure 2.**
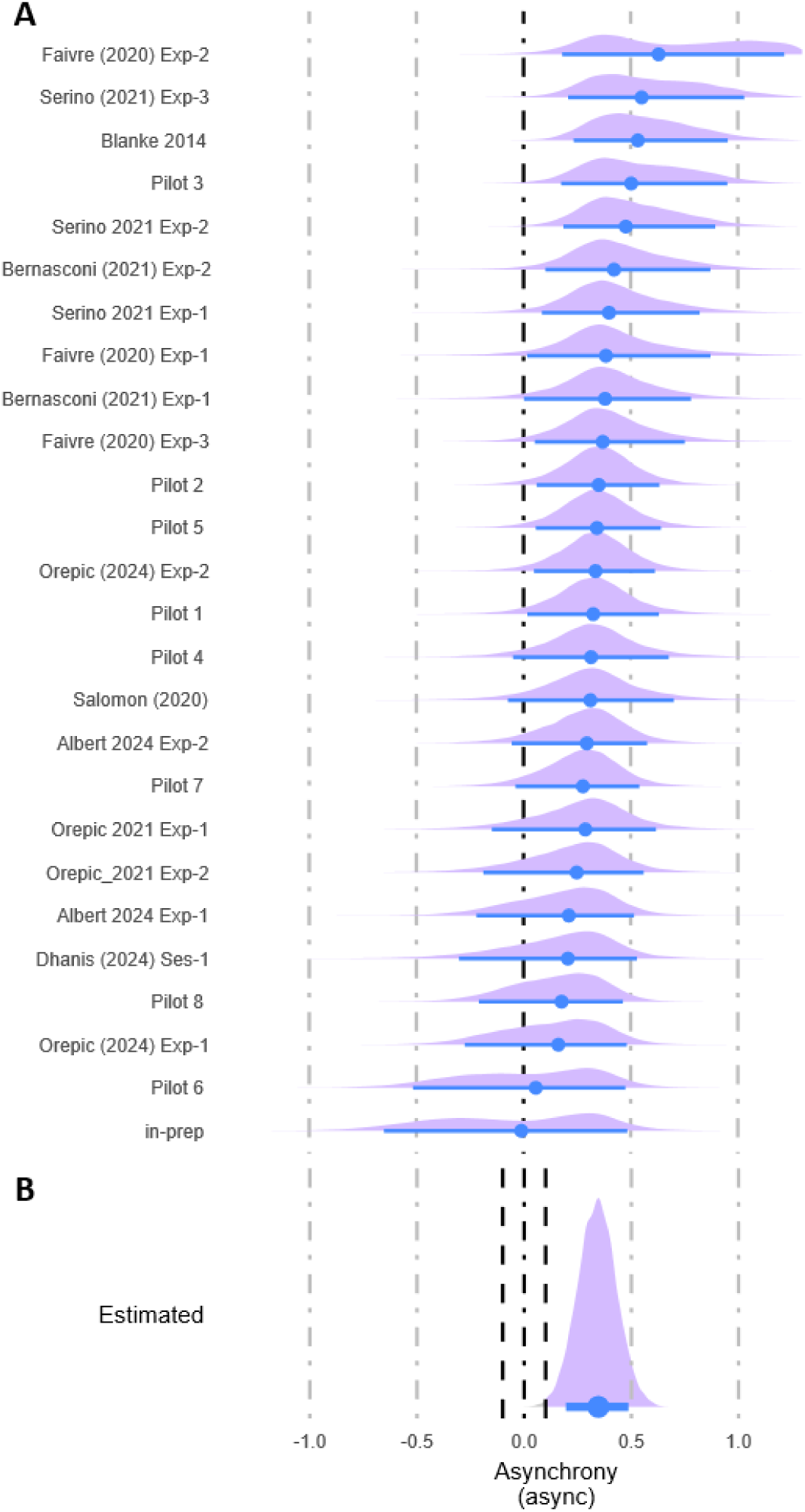
Estimates for the effect of asynchrony across all experiments. **(A)** Forest plot highlighting the different estimates for the asynchrony effect across multiple experiments. The experiments are identified on the left, with more details on Supplementary Table S1. The estimates are shown on the right as cohen’s d, with the distribution of the estimate in purple, the blue circle indicating the mode, and the blue line the 89% high density interval of the estimate. **(B)** Posterior mode estimate for the effect of asynchrony across the experiments. The dashed black lines represent the ROPE.

### 3.3 Effect of demographic and trait characteristics on PH induction

We investigated the effects of age, handedness and delusional ideation on riPH. The latter two were only available for a subset of participants (EHI: 12 experiments, 289 participants, 128 males, mean age of 24.7 years with a SD of 4.9 and range from 17 to 56; PDI: 8 experiments, 178 participants, 64 males, mean age of 24.9 years, with a SD of 5.2 and range from 17 to 56) and special precautions were taken when modelling this data, as described in the methods. Sex at birth was included as a covariate of no interest.

Our results show that delusional ideation positively modulated the ratings of riPH, with higher PDI scores leading to higher riPH ratings (mode: 0.31 SD; HDI: [0.17; 0.43]; Figure 3A). For delusional ideation we further quantified its effect on each specific rating. Overall, participants with higher delusional ideations are more likely to rate the maximum value of “6”, whereas those with lower delusional ideations are more likely to rate “0” (Figure 4). The interaction between asynchrony and PDI did not lead to conclusive results (mode: -0.05 SD, HDI: [-0.21, 0.13], Figure 3B). For both age and handedness, the overlaps between these estimates’ HDIs and the ROPE for the null value were inconclusive on whether they had a reliable effect on riPH (Table 1, Figures 3C-F).

**Figure 3.**
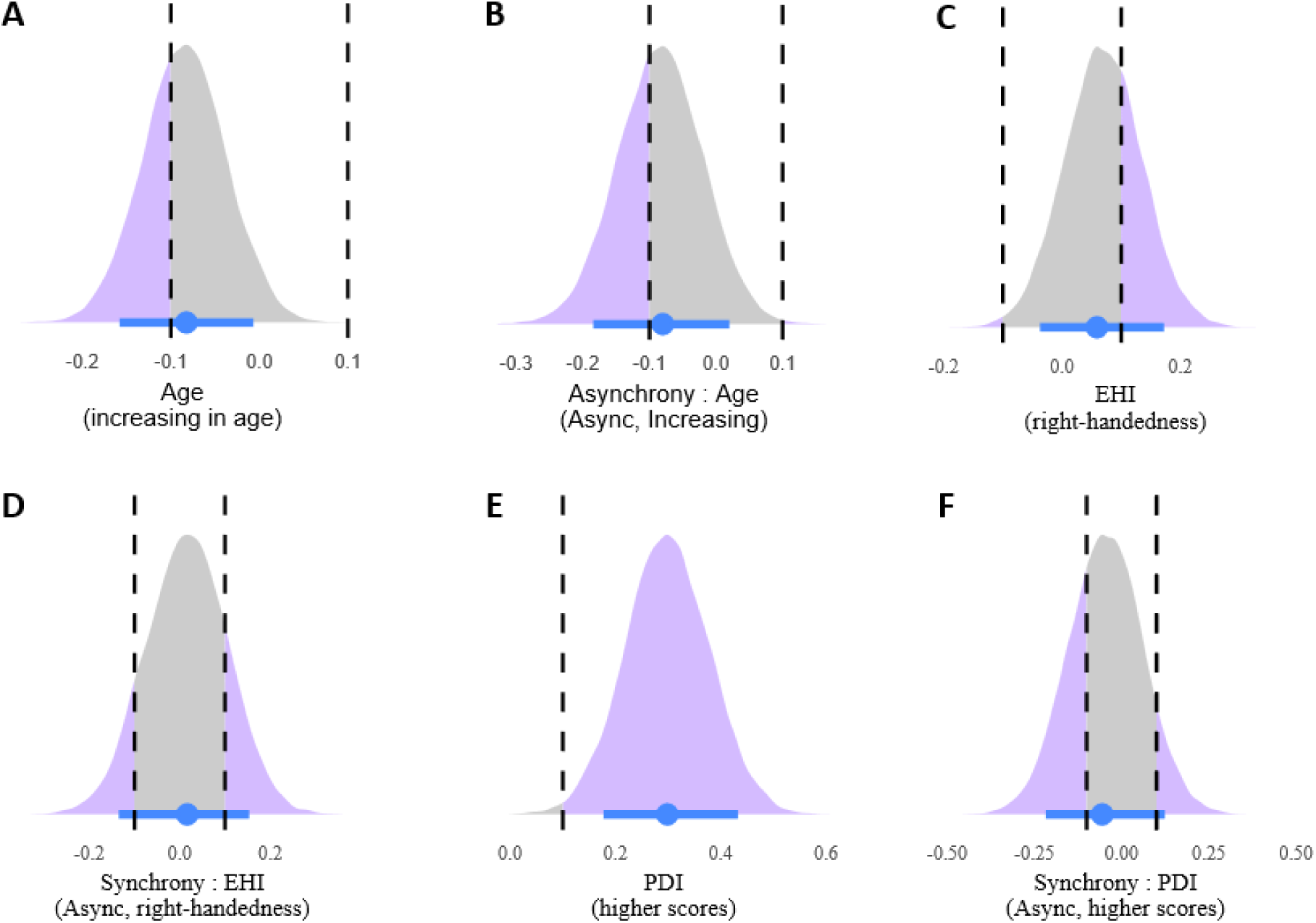
Posterior estimates for age and trait characteristics of participants. Estimates for are shown for characteristics associated with the participants and for their interactions with synchrony. In purple is the distribution of the estimate with the blue circle indicating the mode and the blue line the high density interval (HDI). Vertical dashed black bars, indicate the limits of the region of practical equivalence (ROPE). The distribution of the estimate in show in grey when it overlaps with the ROPE.

**Figure 4.**
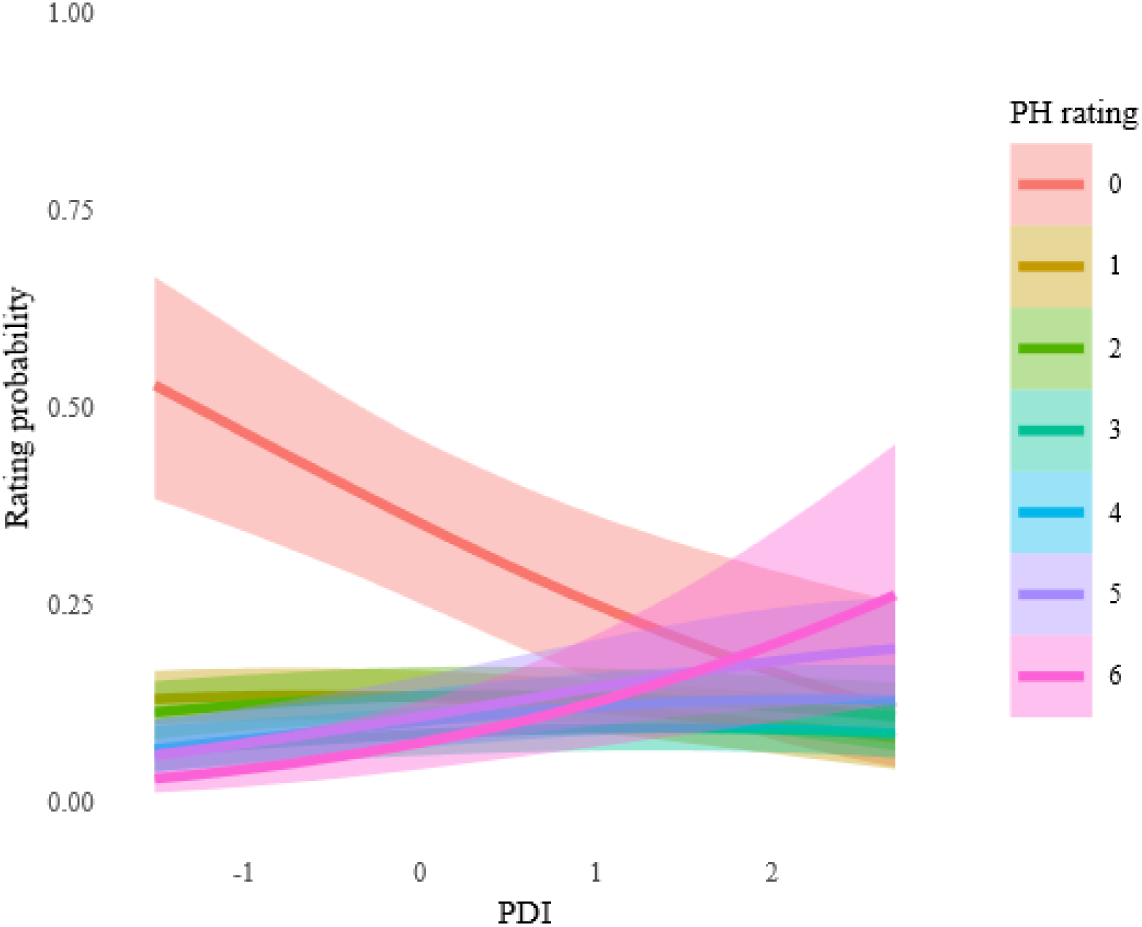
Estimation of the effect of PDI across the ordinal values of the scale. Decomposition of the beta estimates for the effects of POI for each value of the ordinal scale. On the x axis we show the standardize change in the regression coefficient, and on the y axis how the standardized change modulates the probability of rating each value of the scale. For delusional ideations, assessed through POI, a clear distinction is observed for the item "O" which has a high probability of being rated for participants with very low POI, and then sharply decreasing in probability with increasing POI, as compared to item "6" which shows a slight opposite pattern with the probability increasing for individuals with higher delusional ideations. The remaining items remain stable across POI scores.

| Experimental parameter / Demographic trait | Effect size on PH mode standardized | 89% HDI | % of all estimates inside the ROPE | Interaction effect size w/asynchrony | Interaction 89% HDI w/asynchrony | Interaction % all estimates inside the ROPE w/asynchrony |
| --- | --- | --- | --- | --- | --- | --- |
| Asynchrony<br>(effect of asynchrony) | 0.35 | [0.19, 0.48] | < 0.5% | - | - | - |
| Body position<br>(effect of supine) | -0.66 | [-1.10, -0.20] | 2.20% | 0.04 | [-0.27, 0.40] | 35.14% |
| Hand comparator<br>(effect of hand) | -0.61 | [-0.88, -0.33] | < 0.5% | -0.01 | [-0.33, 0.34] | 36.14% |
| Previous exposure<br>(adding previous exposure) | -0.11 | [-0.34, 0.15] | 41.29% | -0.15 | [-0.41, 0.08] | 30.33% |
| Cognitive load<br>(adding cognitive load) | 0.08 | [-0.55, 0.86] | 17.38% | -0.23 | [-0.69, 0.26] | 20.22% |
| Duration<br>(effect of increasing duration) | -0.03 | [-0.28, 0.24] | 46.24% | -0.01 | [-0.19, 0.18] | 61.59% |
| Order<br>(effect of being second) | -0.06 | [-0.26, 0.14] | 52.37% | 0.07 | [-0.16, 0.35] | 41.71% |
| Delusional ideation (PDI)<br>(effect of increasing PDI) | 0.31 | [0.17; 0.43] | 0.71% | -0.05 | [-0.21, 0.13] | 60.50% |
| Age<br>(effect of increasing age) | -0.08 | [-0.16, -0.01] | 63.17% | -0.07 | [-0.19, 0.02] | 59.71% |
| Handedness (EHI)<br>(effect of increasing EHI) | 0.07 | [-0.04, 0.17] | 68.34% | 0.01 | [-0.13, 0.16] | 72.33% |
| Sex at birth<br>(effect of male) | -0.07 | [-0.18, 0.03] | 66.09% | - | - | - |

### 3.4 Effect of different experimental parameters on riPH

With this pooled analysis we also aimed to quantify the effect of different experimental parameters on riPH. Our model revealed that having participants standing or supine changed riPH ratings. PH-induction ratings were lower in both manipulation conditions (asynchronous and synchronous) when using the supine robotic system as compared to the original upright version (mode: -0.65 SD, HDI: [-1.12; -0.15]; Figure 5A). Its effect on asynchrony (i.e., interaction) was inconclusive given the overlap between its HDI and the ROPE for the null value (Figure 5B). Delivering the tactile feedback to the back of the hand instead of the torso also lowered the PH-induction rates in both conditions (mode: - 0.61 SD, HDI: [-0.88, -0.33]; Figure 5C). However, we could not definitively conclude on its interaction effect with asynchrony given the overlap with the ROPE (figure 5D). The estimates for the effect size of previous exposure, cognitive load, duration, and order had significant overlaps between their HDIs and the ROPE for the null value, and consequentially we could not draw definitive conclusions on whether these parameters had an effect or not on riPH (Table 1, Figures 5E-L).

**Figure 5.**
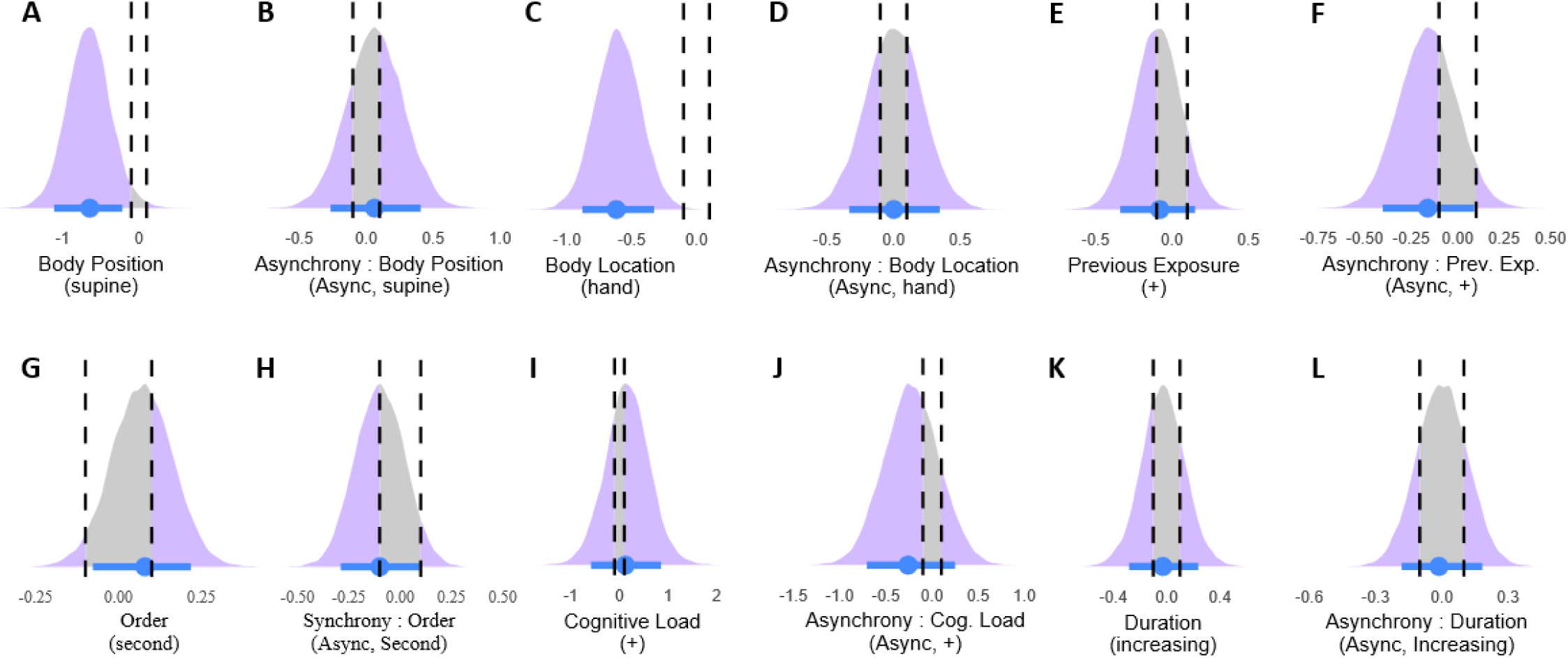
Posterior estimates for the experimental parameters modulating riPH. Posterior estimates for the various experimental parameters or interactions between parameters and synchrony are shown. Below and in parenthesis is indicated the direction of the effect. E.g. Synchrony (async) represents the estimate of the async effect as compared to sync. In purple is the distribution of the estimate with the blue circle indicating the mode and the blue line the high density interval (HDI). Vertical dashed black bars, indicate the limits of the region of practical equivalence (ROPE). The distribution of the estimate in show in grey when it overlaps with the ROPE.

### 3.5 Distribution underlying the riPH rating scale

Given the large amount of data analyzed here, we intended to estimate the distribution underlying 7-point Likert scale used to rate riPH. This was done through the inclusion of flexible threshold parameters for the intercept values of the scale, and visualization of the underlying scale achieved through a generative model that only included these 6 parameters. Figures 6A and 6B show, respectively, the probability of answering each rating of the scale, and how the posteriors sit in relation to a normal distribution (i.e. the underlying distribution of the data). While it is more likely for participants to rate “0” than the remaining values of the scale, this analysis also shows that around 65 to 75% of the 580 participants are sensitive to the riPH procedure, at different levels of the scale. Figure 6C shows how the scale is seen by participants, and figure 6D is distorted to show how participants perceive the distances between points in the ordinal scale (or in other words how they are biased to rate it).

**Figure 6.**
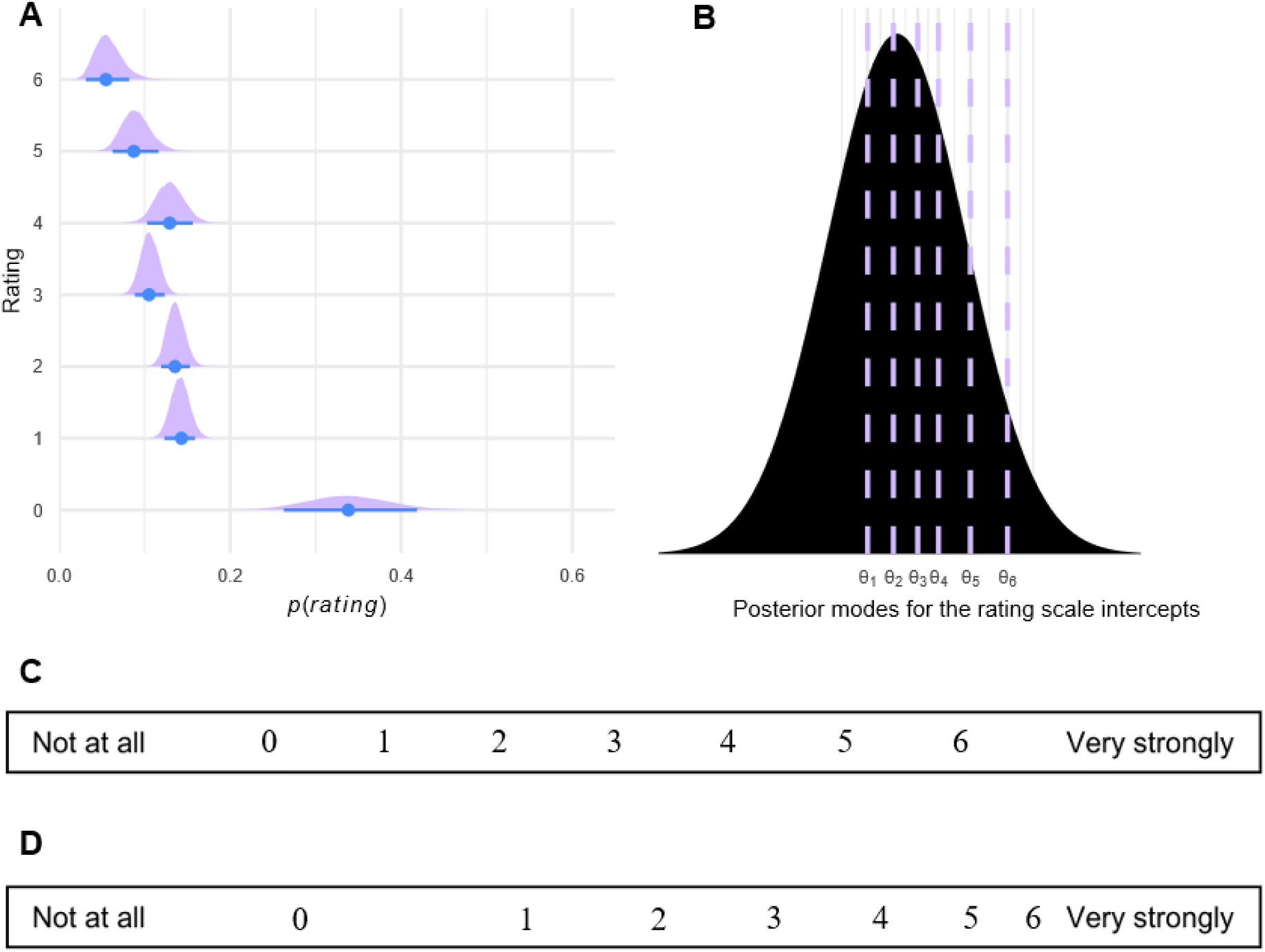
Characteristic of the ordinal rating scale used for riPH. **(A)** Probability of rating each o the items of the rating scale after accounting for the effect of every experimental, demographic, and trait characteristic. In blue is the mode of the probability, with the blue line being the 89% HDI associated to this probability. In purple is the associated distribution. **(B)** Normal distribution with the posterior means of the thresholds separating the values of the ordinal scale. **(C)** Visual representation of the ordinal scale used to rate riPH, as seen visually by the participants. **(D)** The same scale transformed to reflect the inner biases of participants when rating the scale, according to the posterior means described in **(B)**.

In addition to estimating the underlying distribution, we attempted to estimate how much knowing the underlying distribution of the scale could improve the estimation of experimental parameters, including asynchrony in future studies. For this we used *brms* to simulate data based on the properties of the acquired data and then analyzed this data with two types of competing models that either had informed intercept priors or not. According to our estimations on 1000 simulated datasets of experiments with populations varying between 20 and 40 participants, having an informed prior for the intercept improved the estimation of asynchrony in 57% of the cases (judge as estimations being closer to the estimation over all real experiments, and having smaller HDIs).

### 3.6 Effects on passivity experiences, sense of agency and control questions

Here we report the effects of demographic and trait characteristics, and of experimental parameters, on riPE and agency. We note, that whereas riPE induction was investigated for the same number of experiments as riPH, changes in agency were only studied in 9 experiments (189 healthy participants, 80 males, mean age of 24 years-old with a SD of 4.7 and a range of 7 to 37).

Asynchrony modulated the ratings of both riPE (mode = 0.55 SD, HDI = [0.40; 0.68]) and agency (as loss of agency: mode = 0.94 SD, HDI = [0.49, 1.28]). For riPH we had identified that delusional ideation had a positive effect on ratings. Here, we did not see this effect for riPE due to overlaps between its HDI and the ROPE (mode = 0.22 SD, HDI = [0.08, 0.33]), but did observe similar findings for changes in agency, specifically with higher delusional ideation making it more likely for participants to lose agency (mode = 0.39 SD, HDI = [0.21, 0.61]). Regarding the experimental parameters, we had concluded that body position and hand comparator affected the intensity of riPH, but for both riPE and agency we could not conclude on the effect size of these nor of almost all other experimental parameters, due to overlaps between HDIs and the ROPE (Supplementary Tables S4 and S5). The interaction between asynchrony and previous exposure was the only to show an effect, specifically that greater previous exposure decreased the induction effect of asynchrony on riPE (mode: -0.35 SD, HDI: [-0.61, -0.13]; Supplementary Table S4). The illusory feeling of self-touch was also analyzed as an active control question (i.e., an illusion that traditionally works in the synchronous condition). Asynchrony decreased the sensation of self-touch, as expected, and the full results can be seen in Supplementary Table S6.

### 3.7 Relationship between the inductions of PH, PE, and agency

Taking individual participant’s data across all studies, we were interested in assessing the dependencies between riPH, riPE, and agency, as the latter two are typically modulated together with riPH and might influence its experience. For this purpose, we assessed whether the effect of asynchrony on the intensity of riPH (difference in ratings between the asynchronous and synchronous conditions) was modulated by the sensitivity (any positive difference in ratings between the asynchronous and synchronous conditions) to riPE and by loss of agency; and the reverse, i.e., whether the effect of asynchrony on the intensity of riPE was also modulated by sensitivity to riPH and again by loss of agency. In practice this was done, for example for riPH, by analysing model interactions between *sensitivity to changes in agency* and asynchrony, and between *sensitivity to changes in riPE* and asynchrony as well.

#### Mediation analysis for the intensity of riPH

Sensitivity to robotically induced changes in agency did not conclusively modulate the effect of asynchrony on riPH (mode = 0.11 SD, HDI = [-0.42, 0.57], Figure 7A), however, being sensitive to riPE increased asynchrony effect on the intensity of riPH (mode = 0.52 SD, HDI = [0.33, 0.71]; Figure 7B). We did not observe any compounding effect of joint sensitivity to changes in agency and riPE (i.e., triple interaction with asynchrony; mode = 0.22 SD, HDI = [-0.17, 0.67]; Figure 7C).

**Figure 7.**
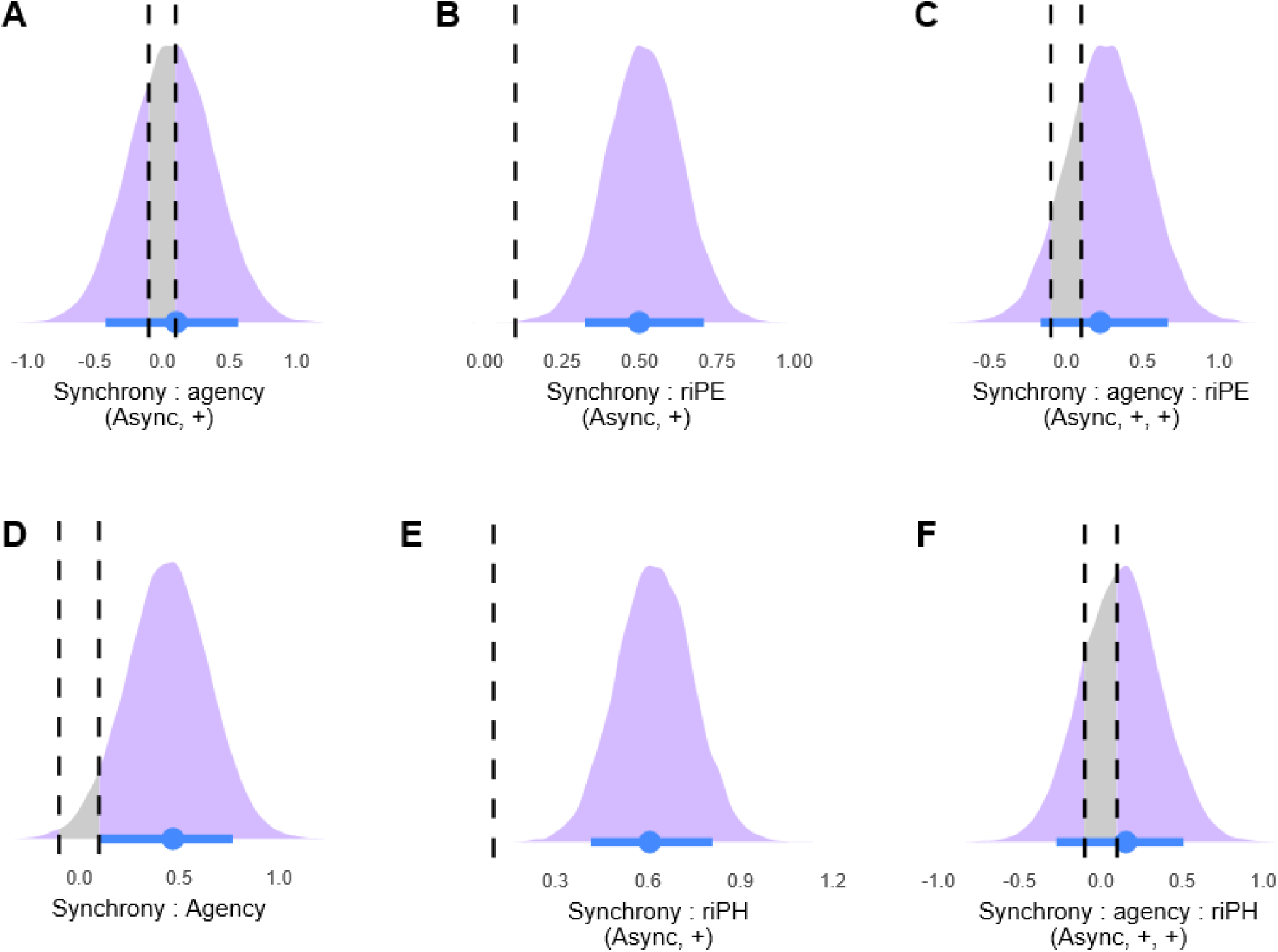
Posterior estimates for the sensitivity mediators. For all posteriors estimates the mode is represented by the blue circle, and the 89% HDI by the associated blue line. Distributional information is in purple, and turns to grey if overlapping with the ROPE (within dashed lines). Mediators for riPH: **(A)** sensitivity to changes in agency, **(B)** sensitivity to riPE, **(C)** both. For riPE: **(D)** sensitivity to changes in agency, **(E)** sensitivity to riPH, **(F)** both.

#### Mediation analysis for the intensity of riPE

Sensitivity to robotically induced changes in agency modulated the asynchrony effect on riPE (mode = 0.33 SD, HDI = [0.12, 0.76]; Figure 7D), and sensitivity to riPH was also shown to increase the asynchrony effect on the intensity of riPE (mode = 0.60 SD, HDI = [0.42, 0.81]; Figure 7E). We did not observe any compounding effect of joint sensitivity to changes in agency and riPH on the intensity of riPE (mode = 0.15 SD, HDI = [-0.27, 0.51]; Figure 7F).

## 4 Discussion

With this analysis of individual-participant data pooled across 26 experiments counting 580 participants, we aimed to investigate multiple aspects of a paradigm which can successfully induce a specific and clinically relevant hallucination, PH, under controlled laboratory conditions. We accurately estimated a medium effect size for the main experimental factor used to induce PH (asynchronous robotic somatomotor stimulation), identified that schizotypal traits measured in healthy participants are associated with higher sensitivity to riPH, determined how body position and other parameters change the intensity of the induction, and supplied statistical priors that should boost the power of future studies. We also dissociated riPH from changes in agency, showing that riPH may not require loss of agency in the way that riPE does, and identified a synergistic relationship between the experiences of riPH and riPE. In sum, we provide a broader understanding of the processes surrounding induction of hallucinations, specifically riPH, as well as important technical details and estimates for future studies investigating riPH.

### 4.1 Relevance of spatial and temporal mechanisms for riPH

The spatiotemporal feature of the robotic somatomotor stimulation has consistently been shown critical for riPH in past studies (Bernasconi et al., 2021; Blanke et al., 2014; Orepic et al., 2021; Salomon et al., 2020; Serino et al., 2021). The current analysis confirms that this temporal conflict, in addition to an existing spatial conflict between movements in the front and touches on the back, leads to an increase in the intensity of riPH. At root of this induction, in the specific context of robotically-mediated somatomotor conflicts, may be a prominent model for motor control in which the somatomotor system uses copies of efferent motor signals to predict the sensory consequences of the motor actions (Kawato, 1999; Miall and Wolpert, 1996; Wolpert et al., 1995; Wolpert and Kawato, 1998). Comparing such predictions to the actual afferent sensory inputs can then inform on whether these are consequence of the motor actions or rather of an external agent (Blakemore et al., 2002; Frith et al., 2000). Key is also that the brain is believed to integrate and interpret multisensory stimuli over a temporal binding window (Colonius and Diederich, 2004; van Wassenhove et al., 2007), which for somatomotor processes shows disintegration between 250 to 500 milliseconds (Kannape et al., 2010; Menzer et al., 2010). Stimuli in this interval can lead to erroneous perceptions (Faivre et al., 2017; Shams et al., 2002). According to this account of motor control (Miall and Wolpert, 1996; Shadmehr et al., 2010; Wolpert et al., 1995), self-other distinction relies on comparing a sensory prediction related to one’s own actions to the actual sensory feedback of such actions. Thus, when there is congruency with the sensorimotor prediction (i.e. in the robotic setup), the actual ascending sensorimotor signals are attenuated, and the performed action is attributed to the self (synchronous manipulation). This, however, differs if the prediction and the ascending sensorimotor signals are incongruent (asynchronous stimulation), leading to reduced attenuation of these incoming signals, and ultimately the action being attributed to an external agent or presence (Heinks-Maldonado et al., 2005; Shergill et al., 2003). We further hypothesize that this process of disrupting motor control, in line with the sensorimotor mismatch hypothesis (Barnby et al., 2023a; Blanke et al., 2014; Case et al., 2020), may mimic what naturally occurs in some patients with PH (Arzy et al., 2006; Blanke et al., 2003; Brugger et al., 1997). In fact, Bernasconi and colleagues presented compelling evidence that this setup mimics many phenomenological characteristics of actual symptomatic hallucination, by identifying a group of Parkinson’s disease patients suffering from symptomatic PH, who were more sensitive to riPH than patients without symptomatic PH (specifically in the asynchronous condition), and who likened the riPH to the symptomatic one experience in daily life (Bernasconi et al., 2021).

### 4.2 The impact of delusional ideations on riPH

We found that higher PDI scores – an indicator of schizotypal traits and the intensity of delusional beliefs – were associated with a general increase in riPH ratings in the tested individuals, meaning that participants with stronger delusional ideation experienced more pronounced riPH during the somatomotor stimulation task. This suggests that schizotypal traits, such as heightened perceptual sensitivity and a propensity for exacerbated cognitive biases, likely contributed to the experience of riPH. These observations are in-line with previous research from other domains showing: that individuals with schizotypal traits are more prone to auditory and visual hallucinations (de Leede-Smith and Barkus, 2013; Debbané et al., 2014; Dolphin et al., 2015); that proxies for auditory-verbal hallucinations such as false alarms in an auditory detection task, are increased in healthy individuals scoring higher on the PDI (Orepic et al., 2024); and that healthy individuals with paranoid ideations and delusional proneness are more susceptible to induction of somatomotor changes (Germine et al., 2013; Kállai et al., 2015).

One hypothesis to explain these observations is that errors in predictive coding play a role in hallucination induction. According to the Bayesian model of perception (Fletcher and Frith, 2009; Mirza et al., 2019), top-down processing likely amplifies ambiguous stimuli into hallucinatory experiences. Individuals may prioritize internal expectations over sensory input, leading to the misinterpretation or unnecessary significance being attributed to ambiguous stimuli, such as the somatomotor conflicts experienced during the robotic task. This can result in distorted perceptions (Corlett et al., 2019; Schmack et al., 2013) and over time, these distortions may further reinforce delusional ideation by shifting cognitive biases (i.e., prior beliefs) (Willard and Norenzayan, 2013). Specifically, the brain’s reliance on prior knowledge, expectations, and beliefs, to interpret unclear somatosensory input, may generate aberrant perceptual phenomena such as the feeling of someone’s presence (Fletcher and Frith, 2009). However, the increased sensitivity to riPH in participants with higher PDI scores – indicative of schizotypal traits – may also arise from changes in their sense of agency during the somatomotor task (Asai et al., 2016; Farrer and Frith, 2002; Lallart et al., 2009). Individuals with high schizotypy incur in attenuation deficits of self-produced motor-tactile stimuli (Asimakidou et al., 2022) and may struggle to distinguish between internally generated and external stimuli. This was evidenced in our results, where participants reported higher passivity experiences and loss of agency in the asynchronous condition, with the latter being further exacerbated by higher schizotypy. Previous studies have shown that passivity experiences can arise from a disrupted sense of agency in both healthy individuals (Orepic et al., 2020) and those with schizophrenia spectrum disorders, contributing to hallucinations and delusional ideas (Graham-Schmidt et al., 2018; Stripeikyte et al., 2021). This lack of clarity regarding the origin of perception can increase the likelihood of hallucinations, as individuals misattribute sensory experiences, leading to experiences resembling delusions of control and a sense that their actions are externally influenced. In sum, our findings highlight how schizotypal traits can influence the boundary between normal perception, belief systems, and proneness to hallucinations, and while the above hypotheses are not mutually exclusive, the current analysis of pooled data across multiple experiments allowed us to further investigate the relationship between riPH, agency and passivity.

### 4.3 Robot-induced PH, passivity experiences, and agency

To study the potential relationship between riPE and riPH, as well as their potential dependencies on agency, we first established that the robotic task was indeed leading to changes in the sense of agency, expressed as loss of agency in the asynchronous condition. This a common finding in experiments that rely on asynchronous somatomotor tasks, albeit generally focused on hand movements (for a meta-analysis on motor control and agency: Zito et al., 2020). Having guaranteed this, we then investigated via mediation analyses whether riPE and riPH had dependencies on the sense of agency, and determined that being prone to losing agency boosted the experience of riPE in the asynchronous condition. Agency and passivity experiences are both more easily manipulated in healthy individuals with higher delusional ideations (Louzolo et al., 2015), and clinical research shows that PE depends on agency deficits in schizophrenic patients (Moore, 2016), supporting the facilitating role of loss of agency on riPE observed here. However, for riPH we could not establish that the same changes in agency contributed in any way to its perception. These distinct results suggest that riPH does not rely as strongly in loss of agency as riPE, and that by extension it must also rely on partly distinct brain mechanism.

To further investigate the relationship between riPE and riPH, we then analyzed whether being sensitive to riPE (in addition to sensitivity to changes in agency) boosted the intensity of riPH, and vice-versa. On this, the mediation analyses supported that the sensitivity to either experience exacerbated the effect of asynchrony making the intensity of the other experience even stronger (i.e., being sensitive to riPH led to a more intense riPE with asynchrony, and vice-versa), indicating that the two share at least a partial synergism between them. Coupling the two results, we identified that behaviorally, while only riPE is boosted by loss of agency, the experiences of riPE and riPH are mutually increased by one another. Hence, it is likely that the two also rely on both common and distinct brain mechanisms. Recent neuroimaging data gives strength to this account, as two distinct processes of communication between brain networks were identified for riPH and riPE, yet the brain process underlying riPE could already be partially observed for riPH (albeit not in a sufficient manner to explain riPH; Dhanis et al., 2022). More work is needed to reveal the common and distinct behavioral and brain mechanisms of riPH, riPE and sense of agency. In sum, the present analysis supports a synergistic relationship between riPH and riPE and a distinct dependency of each of these with the sense of agency. However, these dependencies may not be the only factors that distinguish riPH from riPE as these were not equally affected by the same experimental parameters (as discussed further below).

### 4.4 The effect of experimental parameters and statistical priors

The original standing robot (Blanke et al., 2014) was shown to induce a generally stronger riPH than its supine MR-compatible counterpart (Bernasconi et al., 2021; Hara et al., 2014). Several reasons might have led to this effect. First, in the supine position participants have and feel the MRI bed immediately below their back with no physical space behind/below them (i.e., for a “presence”). However, this may only be partly true. In the standing robot setup it could be argued that the robot behind the participant also interferes with the space that would be available for a presence, yet this has not hindered induction in healthy individuals (Blanke et al., 2014; Serino et al., 2021) nor in patients (Bernasconi et al., 2021). In addition, PH has been induced through focal electrical stimulation to the brain, in a patient who was in supine position, and the presence still occurred below or partially intermingled with the bed (Arzy et al., 2006). When taking these factors into consideration, the participant’s notion of perceived space should not be sufficient to explain this difference. Another explanation has to do with technical differences between the two robots. The standing robot allows for more degrees of freedom in the movements performed by the participants and allows for greater control over the pressure exerted by the back-robot, as compared to the MR-compatible robot (i.e., closer mimicking of the movement and force). This hypothesis aligns with the importance of somatomotor processes for riPH (i.e., sensorimotor mismatch hypothesis), however further studies focusing directly on these technical components would be beneficial to confirm this. Notwithstanding, despite a decrease in general intensity, no interaction between the predictors of asynchrony and body-position were observed, suggesting that the effect of asynchrony is preserved and riPH is still achieved in supine position. Experimentally, researchers should keep in mind this general decrease in intensity and attempt to increase their population size to sustain adequate statistical power.

Providing somatosensory feedback to the hand as compared to the back also produced a weaker riPH in both manipulation conditions (synchronous, asynchronous). This is compatible with proposals that the brain’s torso representation is critical for multisensory and somatomotor processes that are of relevance for global and unitary aspects of bodily self-consciousness (Blanke, 2012; Blanke et al., 2015; Park and Blanke, 2019). Different studies show that trunk stimulation can disrupt such global aspects of bodily self-consciousness such as self-identification, self-location, and first-person perspective (Ehrsson, 2007; Ionta et al., 2011; Lenggenhager et al., 2007; Petkova et al., 2011). Moreover, neurological damage affecting these global body representation has been found to lead to autoscopic phenomena, including out-of-body experiences (Ionta et al., 2011), heautoscopy (Heydrich and Blanke, 2013), or PH (Arzy et al., 2006; Blanke et al., 2014). Conversely representations of the hand and fingers activate distinct neurons and brain regions in posterior parietal cortex and premotor cortex (Blanke et al., 2015) and are related to isolated or part-to-global aspects of bodily self-consciousness such as hand, foot, or finger ownership (Botvinick and Cohen, 1998; Ehrsson et al., 2004; Pozeg et al., 2015). Combined with the present results on tactile feedback location (hand vs trunk) this suggests that trunk-centered feedback (vs. hand feedback) leads to stronger riPH, because it interferes with participants’ global body representation, giving not just rise to riPE and loss of agency, but also the perception of another person, riPH.

Of the other parameters analyzed, we were not able to assert definitively whether they had a positive, negative, or null effect on riPH. Previous exposure, order, duration, age, and handedness, all had estimated effects sitting well within the ROPE but whose tails extended beyond it and accordingly we are more confident that these parameters do not have an effect on riPH. In terms of experimental control, this is particularly relevant for duration and previous exposure, as it potentially allows researchers to do longer experiments and repeat them over time, without compromising the integrity of PH induction. Interestingly, while this may be true for riPH, we did identify that previous exposure to the robot reduces the induction effect of asynchrony on riPE. Together this may mean that riPH is more robust over repeated sessions than riPE, potentially related to increased reliance of riPH on the body distortions created by our setup (described above), as compared to riPE, or for example on the distinct neural mechanisms underpinning each induced sensation (Dhanis et al., 2022). Future work is be needed to elucidate this. Lastly, an exception to the apparent no-effect is cognitive load. The posterior distribution for the effect size of this parameter was larger than the others, meaning that the uncertainty associated with is higher. Hence, it could be possible that cognitive load has a positive or negative effect on induction. For example, it might be that higher intensities of cognitive load have an effect, but lower intensities do not. However, one study investigating riPH has looked specifically at this effect using a N-back task, and found no effect of cognitive load (Serino et al., 2021), supporting the hypothesis that cognitive load does not affect riPH. Notwithstanding, we still advise researchers to approach this with care.

Finally, thanks to the statistical tools used here, we were also able to estimate the effect sizes and HDIs of the experimental parameters as well as those of the intercepts of the rating scale for riPH. We have briefly shown that prior knowledge of these intercepts boosts the statistical power in simulations with the typical number of participants used in these experiments. To improve future studies, we have made all these priors available here: https://github.com/HDhanis/pooledPH.

### 4.5 Limitations

With this work providing important contributions to the study of riPH, some limitations must also be noted. First, because all data derived from a single laboratory, it lacks independent multi-site replication, and external generalizability should ideally be tested in future multi-lab studies. Nonetheless, our findings still reflect extensive internal replication, consistency across multiple setups with experimental changes, and inference of effect sizes in a large sample size. Second, it is important to stress that although convergency of our models was verified and ensured, the distributions of some experimental parameters were large with respect to their expected effect sizes, limiting the usefulness of the extracted information. Third, we made binary simplifications for the presence or not of cognitive load and previous exposure, as complexity did not allow us to model their respective intensities and durations. Hence, it is possible that the degree to which these parameters varied when present in the experiments, did model riPH in a way not captured here (e.g., a concomitant task with minor cognitive load might not impact riPH but one with major cognitive load might to do so in either direction). Finally, regarding the mediation analysis, our initial hypothesis is not directly testable as a two-step mediation procedure and requires a temporal structure to disambiguate the order of mediation. This is unfortunately not accessible in these types of experiments. We nonetheless used an approach that to an extent encodes but does not enforce such a relationship.

## 5 Conclusion

In brief, with this analysis that pooled data from all our PH-induction experiments we furthered the knowledge of processes surrounding the induction of hallucinations such as riPH, identified a segment of the general population that is more sensitive to its induction due to higher delusional ideations, and quantified the effects of induction and other experimental parameters related to the somatomotor task. In addition, we extended the results of this analysis and neuroimaging studies (Dhanis et al., 2022) on the relationship between riPH and two other important manipulated somatomotor processes: agency and passivity. Specifically, we showed that riPH does not seem to depend on changes in agency as passivity does, but that there is still a synergistic relationship between riPH and passivity. We based these contributions on strong statistical tools, which expanded our understanding of hallucination induction and provide now useful priors for future studies assessing riPH. Furthermore, all these results, even if in healthy individuals, are highly informative for a setup that is now being used in various clinical applications, including in Parkinson’s disease (Bernasconi et al., 2021).

