## Supplementary Info for "Presence Hallucination Induction through Robotically Mediated Somatomotor Conflicts – a pooled analysis of 26 experiments"

### Supplementary Information

**Supplementary Table S1** List of included studies with respective experiments. The experiment ID corresponds to the ID used in the released excel data file.

| Experiment ID | Study and experiment | Number of participants |
|---|---|---|
| **1** | Pilot 1 by Dhanis, H. and Bernasconi, F. | 12 |
| **2** | Pilot 2 by Dhanis, H. and Bernasconi, F. | 23 |
| **3** | Pilot 3 by Bernasconi, F. | 26 |
| **4** | Bernasconi et al. (2021) *Science Translational Medicine*: Mock fMRI | 24 |
| **5** | Bernasconi et al. (2021) *Science Translational Medicine*: fMRI | 25 |
| **6** | Pilot 4 by Stripeikyte, G. | 24 |
| **7** | Orepic et al., (2021) *Schizophrenia Research*: Experiment 1 | 30 |
| **8** | Orepic, et al. (2021) *Schizophrenia Research*: Experiment 2 | 30 |
| **9** | Pilot 5 by Potheegadoo, J. | 26 |
| **10** | Pilot 6 by Potheegadoo, J., and Blondiaux, E. | 15 |
| **11** | Serino et al. (2021) *iScience*: Experiment 1 | 35 |
| **12** | Serino et al. (2021) *iScience*: Experiment 3 | 19 |
| **13** | Blanke et al. (2014) *Current Biology*: Study 3 | 21 |
| **14** | Pilot 7 by Orepic, P. | 15 |
| **15** | Salomon et al. (2020) *Schizophrenia Bulletin*: Control participants | 20 |
| **16** | Serino et al. (2021) *iScience*: Experiment 4 | 21 |
| **17** | Orepic et al. (2024) *Psychological Medicine*: Experiment 1 | 24 |
| **18** | Faivre et al. (2020) *Cortex*: Experiment 1 | 18 |
| **19** | Faivre et al. (2020) *Cortex*: Experiment 2 | 19 |
| **20** | Faivre et al. (2020) *Cortex*: Experiment 3 | 18 |
| **21** | Orepic et al. (2024) *Psychological Medicine*: Experiment 2 | 24 |
| **22** | Pilot 8 by Franza, M. | 24 |
| **23** | Dhanis, H., et al. (2024) *Communications Biology* | 20 |
| **24** | Pilot 9 by Franza, M. | 17 |
| **25** | Albert et al. (2024) *Nature Communications* | 25 |
| **26** | Albert et al. (2024) *Nature Communications* | 25 |

**Supplementary Table S2** All questions collected for all experiments. Question ID refers to the ID used in the excel data provided. Please note that the phrasing of some questions slightly changed throughout the 8 years in which these experiments have been collected, however, they were monitored and assessed by an experienced neuropsychologist to guarantee that the valence and what they assess remained the same. We group questions with very similar phrasing and that assessed the same thing, under the same question.

| Question ID | Question | Aspect measured |
|---|---|---|
| **1** | I felt as if I had no body | Control |
| **2** | I felt as if I was touching my body | Self-touch |
| **3** | I felt as if I was touching someone else's body | Touching other |
| **4** | I felt as if I was behind my body | Self-location |
| **5** | I felt as if I had more than one body | Control |
| **6** | I felt as if someone else was touching my body | Passivity experiences |
| **7** | I felt as if someone was standing close to me (next to me or behind me) | Presence Hallucination |
| **8** | I felt as if someone was standing in front of my body | Presence Hallucination (at front – used for hand experiments) |
| **9** | I felt as if I had two right hands | Control |
| **10** | I felt as if there was a kind/gentle presence behind me | Presence Hallucination: valence follow-up *(only in few experiments)* |
| **11** | I felt as if there was a weird/unpleasant presence next to me | Presence Hallucination: valence follow-up *(only in few experiments)* |
| **12** | I felt as if I could hear someone else's voice in my mind | Thought insertion |
| **13** | I felt as if I could share my thoughts with someone else | Thought broadcasting |
| **14** | I had the impression I could not control my own thoughts | Thought agency |
| **15** | I felt as if someone could read my mind or hear my thoughts | Thought intrusion |
| **16** | I felt as if I was not controlling my movements or actions | Sense of Agency (as loss of agency) |
| **17** | I felt anxious or stressed | Anxiety |
| **18** | I felt as if I was in front of my body | Self-location |
| **19** | Presence position (0-no answer, 1-left, 2-middle, 3-right, 4-other) | Presence Hallucination: follow-up *(only in few experiments)* |
| **20** | I felt as if I was touched by a robot | Control |
| **21** | I felt as if someone else was controlling my movements or actions | Alien agent |

**Supplementary Table S3** Experimental parameters, trait characteristics and SoA question for each experiment

| Experiment | Duration (seconds) | Body Position | Hand Comparator | Previous Exposure | Cognitive Load | Has PDI | Has EHI | Has Loss of Agency question |
|---|---|---|---|---|---|---|---|---|
| **Pilot 1 – Dhanis, H. and Bernasconi, F.** | 120 | Up-right | Yes | No | No | No | No | No |
| **Pilot 2 – Dhanis, H. and Bernasconi, F.** | 120 | Up-right | Yes | No | No | No | No | No |
| **Pilot 3 – Bernasconi, F.** | 120 | Up-right | No | No | No | No | No | No |
| **Bernasconi et al. (2021) Science Translational Medicine: Mock fMRI** | 180 | Supine | No | Yes | No | Yes | Yes | Yes |
| **Bernasconi et al. (2021) Science Translational Medicine: fMRI** | 30 | Supine | No | Yes | No | No | Yes | No |
| **Pilot 4 – Stripeikyte, G.** | 120 | Supine | No | Yes | No | No | Yes | No |
| **Orepic et al., (2021) Schizophrenia Research: Experiment 1** | 270 | Up-right | No | Yes | Yes | Yes | Yes | No |
| **Orepic, et al. (2021) Schizophrenia Research: Experiment 2** | 270 | Up-right | Yes | Yes | Yes | Yes | Yes | No |
| **Pilot 5 – Potheegadoo, J.** | 180 | Up-right | No | No | No | Yes | Yes | No |
| **Pilot 6 – Potheegado, J. And Blondiaux, E.** | 120 | Up-right | No | No | No | No | No | Yes |
| **Serino et al. (2021) iScience: Experiment 1** | 300 | Up-right | No | No | Yes | No | No | No |
| **Serino et al. (2021) iScience: Experiment 3** | 60 | Up-right | No | No | No | No | No | No |
| **Blanke et al. (2014) Current Biology: Study 3** | 180 | Up-right | No | No | No | No | No | No |
| **Pilot 7 – Orepic, P.** | 120 | Up-right | No | No | No | Yes | Yes | Yes |
| **Salomon et al. (2020) Schizophrenia Bulletin: Control participants** | 180 | Up-right | No | Mixed (some yes others no) | No | No | No | No |
| **Serino et al. (2021) iScience: Experiment 4** | 60 | Up-right | No | Mixed (some yes others no) | No | No | Yes | No |
| **Orepic et al. (2024) Psychological Medicine: Experiment 1** | 120 | Up-right | No | No | No | Yes | Yes | No |
| **Faivre et al. (2020) Cortex: Experiment 1** | 500 | Up-right | No | No | Yes | No | No | No |

| | | | | | | | | |
|---|---|---|---|---|---|---|---|---|
| **Faivre et al. (2020) Cortex: Experiment 2** | 60 | Up-right | No | Yes | No | No | No | No |
| **Faivre et al. (2020) Cortex: Experiment 3** | 60 | Up-right | Yes | Yes | No | No | No | No |
| **Orepic et al. (2024) Psychological Medicine: Experiment 2** | 120 | Up-right | No | No | No | Yes | No | Yes |
| **Pilot 8 – Franza, M.** | 120 | Up-right | No | Mixed (some yes others no) | No | No | No | Yes |
| **Dhanis, H., et al. (2024) Communications Biology: Day 1** | 30 | Supine | No | No | No | No | Yes | Yes |
| **Pilot 9 – Franza, M.** | 120 | Up-right | Yes | Mixed (some yes others no) | No | Yes | No | Yes |
| **Albert et al. (2024) Nature Communications** | 60 | Up-right | No | No | No | No | Yes | Yes |
| **Albert et al. (2024) Nature Communications** | 60 | Up-right | No | No | No | No | Yes | Yes |

**Supplementary Table S4** Effect of experimental parameters on the induction of robot induced Passivity Experiences

| Experimental parameter / Demographic trait | Effect size on PH mode standardized | 89% HDI | % of all estimates inside the ROPE | Interaction effect size w/asynchrony | Interaction 89% HDI w/asynchrony | Interaction % all estimates inside the ROPE w/asynchrony |
|---|---|---|---|---|---|---|
| **Asynchrony** (effect of asynchrony) | 0.55 | [0.40, 0.68] | 0 | - | - | - |
| **Body position** (effect of supine) | -0.28 | [-0.67, 0.19] | 19.50% | 0.08 | [-0.22, 0.41] | 35.20% |
| **Hand comparator** (effect of hand) | -0.32 | [-0.58, -0.07] | 9.50% | -0.18 | [-0.50, 0.12] | 24.66% |
| **Previous exposure** (adding previous exposure) | 0.07 | [-0.14, 0.33] | 44.19% | -0.35 | [-0.61, -0.13] | 3.67% |
| **Cognitive load** (adding cognitive load) | -0.20 | [-0.86, 0.55] | 16.51% | 0.19 | [-0.29, 0.63] | 23.23% |
| **Duration** (effect of increasing duration) | 0.07 | [-0.19, 0.32] | 44.71% | -0.08 | [-0.27, 0.09] | 49.06% |
| **Order** (effect of being second) | -0.05 | [-0.18, 0.10] | 68.95% | 0.13 | [-0.06, 0.31] | 38.96% |
| **Delusional ideation (PDI)** (effect of increasing PDI) | 0.22 | [0.08, 0.33] | 7.26% | 0.08 | [-0.10, 0.24] | 53.99% |
| **Age** (effect of increasing age) | -0.16 | [-0.24, 0.09] | 7.58% | -0.01 | [-0.12, 0.08] | 87.94% |
| **Handedness (EHI)** (effect of increasing EHI) | -0.10 | [-0.19, 0.01] | 53.61% | 0.06 | [-0.09, 0.19] | 68.02% |
| **Sex at birth** (effect of male) | -0.10 | [-0.19, 0.00] | 53.23% | - | - | - |

**Supplementary Table S5** Effect of experimental parameters on changes in Sense of Agency

| Experimental parameter / Demographic trait | Effect size on PH mode standardized | 89% HDI | % of all estimates inside the ROPE | Interaction effect size w/asynchrony | Interaction 89% HDI w/asynchrony | Interaction % all estimates inside the ROPE w/asynchrony |
|---|---|---|---|---|---|---|
| **Asynchrony** (effect of asynchrony) | 0.94 | [0.49, 1.28] | < 0.5% | - | - | - |
| **Body position** (effect of supine) | 0.33 | [-0.58, 1.32] | 11.40% | -0.50 | [-1.00, -0.04] | 6.98% |
| **Hand comparator** (effect of hand) | 0.54 | [0.02, 1.13] | 6.03% | -0.70 | [-1.36, -0.04] | 4.98% |
| **Previous exposure** (adding previous exposure) | 0.06 | [-0.41, 0.59] | 24.00% | 0.19 | [-0.41, 0.80] | 17.97% |
| **Cognitive load** (adding cognitive load) | n/a | n/a | n/a | n/a | n/a | n/a |
| **Duration** (effect of increasing duration) | -0.40 | [-1.37, 0.46] | 10.04% | 0.04 | [-0.47, 0.55] | 24.06% |
| **Order** (effect of being second) | 0.07 | [-0.20, 0.36] | 39.15% | -0.34 | [-0.69, 0.06] | 13.04% |
| **Delusional ideation (PDI)** (effect of increasing PDI) | 0.39 | [0.21, 0.61] | 0.65% | -0.22 | [-0.46, 0.71] | 23.88% |
| **Age** (effect of increasing age) | -0.17 | [-0.34, -0.03] | 20.36% | 0.11 | [-0.08, 0.30] | 42.17% |
| **Handedness (EHI)** (effect of increasing EHI) | -0.08 | [-0.28, 0.11] | 49.43% | 0.01 | [-0.28, 0.25] | 44.71% |
| **Sex at birth** (effect of male) | -0.20 | [-0.37, 0.00] | 19.19% | - | - | - |

**Supplementary Table S6** Effect of experimental parameters on self-touch

| Experimental parameter / Demographic trait | Effect size on PH mode standardized | 89% HDI | % of all estimates inside the ROPE | Interaction effect size w/asynchrony | Interaction 89% HDI w/asynchrony | Interaction % all estimates inside the ROPE w/asynchrony |
|---|---|---|---|---|---|---|
| **Asynchrony** (effect of asynchrony) | -0.62 | [-0.76, -0.47] | 0% | - | - | - |
| **Body position** (effect of supine) | 0.14 | [-0.33, 0.60] | 24.50% | 0.14 | [-0.18, 0.49] | 29.08% |
| **Hand comparator** (effect of hand) | -0.14 | [-0.39, 0.12] | 34.93% | 0.36 | [0.04, 0.66] | 9.70% |
| **Previous exposure** (adding previous exposure) | 0.01 | [-0.23, 0.24] | 50.82% | 0.01 | [-0.23, 0.24] | 23.07% |
| **Cognitive load** (adding cognitive load) | -0.52 | [-1.28, 0.12] | 7.66% | 0.22 | [-0.22, 0.70] | 19.16% |
| **Duration** (effect of increasing duration) | 0.13 | [-0.11, 0.40] | 32.98% | -0.07 | [-0.26, 0.10] | 51.76% |
| **Order** (effect of being second) | 0.17 | [-0.01, 0.35] | 21.94% | -0.25 | [-0.47, 0.01] | 12.65% |
| **Delusional ideation (PDI)** (effect of increasing PDI) | -0.01 | [-0.15, 0.15] | 85.46% | -0.09 | [-0.30, 0.11] | 49.06% |
| **Age** (effect of increasing age) | -0.09 | [-0.16, -0.01] | 62.20% | 0.05 | [-0.05, 0.16] | 74.79% |
| **Handedness (EHI)** (effect of increasing EHI) | 0.01 | [-0.13, 0.13] | 92.49% | -0.07 | [-0.24, 0.10] | 63.15% |
| **Sex at birth** (effect of male) | 0.09 | [-0.01, 0.19] | 55.74% | - | - | - |

### Diagnostic Plots

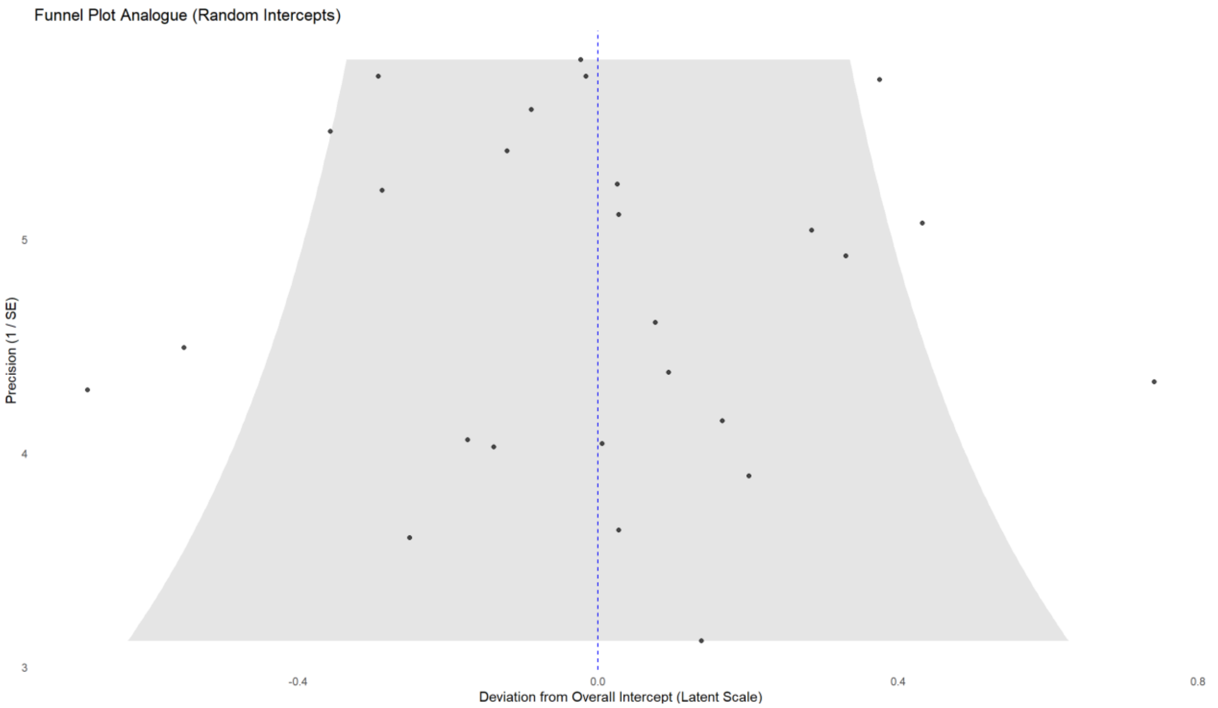


Supplementary Figure S1: Funnel plot analogue for an ordinal model with flexible thresholds showing study-specific random intercept estimates. Each point represents a study's deviation from the overall intercept on the latent scale, plotted against its precision (inverse of the standard error). The shaded grey area indicates the expected 95% confidence region under the null hypothesis of no between-study heterogeneity. Points lying outside this area may suggest outliers or greater heterogeneity across studies. This was limited in our case. The dashed vertical line at zero corresponds to no deviation from the average intercept.

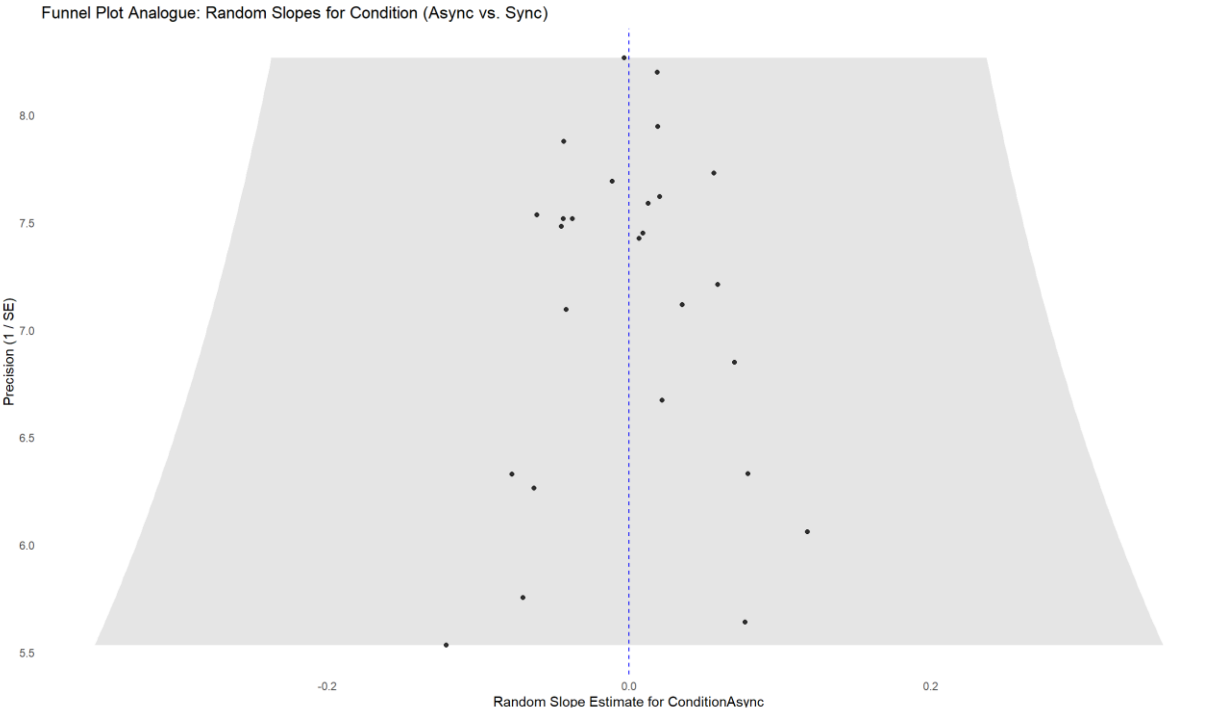


Supplementary Figure S2: Funnel plot analogue displaying study-specific random slope estimates for the effect of Asynchrony as relative to Synchrony. Each point represents a study's deviation in the condition effect, plotted against its precision (1 / standard error). The shaded grey area represents the expected 95% confidence region assuming no small-study bias or heterogeneity. The dashed vertical line at zero corresponds to no deviation from the average condition effect. Studies falling outside the shaded region may indicate potential outliers or heterogeneity.

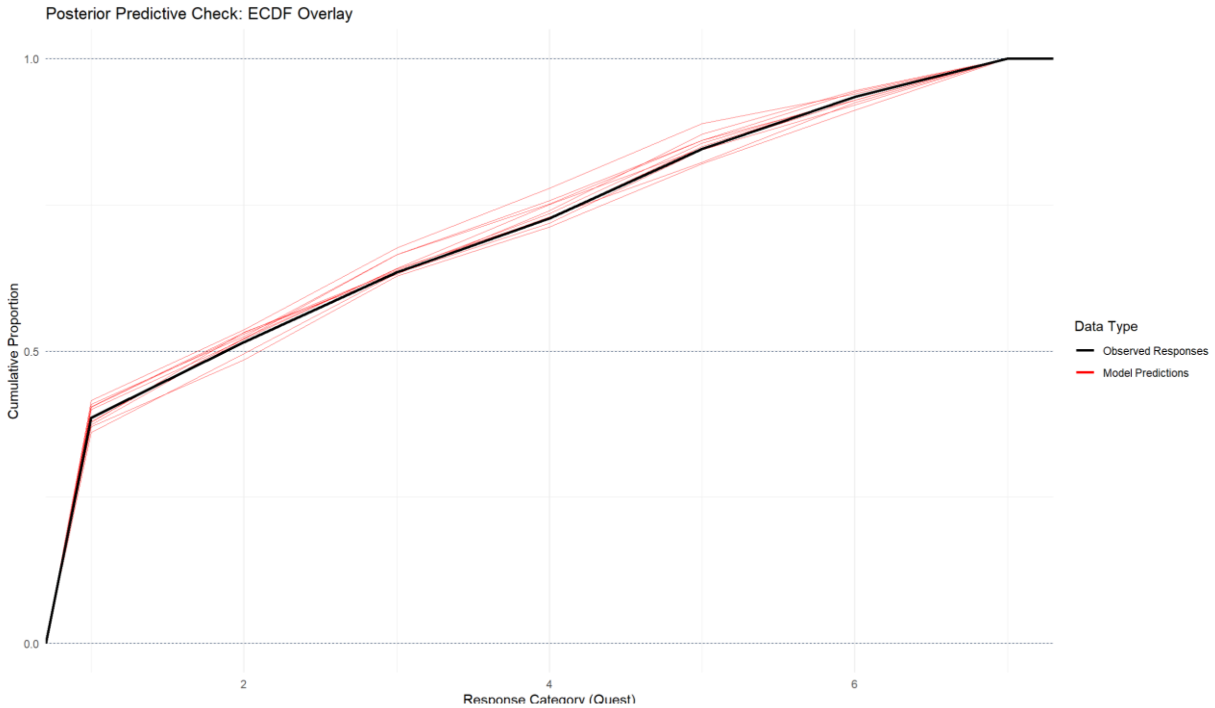


Supplementary Figure S3: Posterior predictive check using empirical cumulative distribution functions (ECDFs) for the ordinal outcome variable. The black line represents the empirical distribution of the observed responses, while the red lines show replicated data drawn from the posterior predictive distribution (n = 10 draws). Overlap between observed and replicated distributions indicates that the model captures the overall distributional shape of the outcome well.
